# Cre/*lox*-controlled spatio-temporal perturbation of FGF signaling in zebrafish

**DOI:** 10.1101/302174

**Authors:** Lucia Kirchgeorg, Anastasia Felker, Elena Chiavacci, Christian Mosimann

## Abstract

Fibroblast Growth Factor (FGF) signaling guides multiple developmental processes including body axis formation and specific cell fate patterning. In zebrafish, genetic mutants and chemical perturbations affecting FGF signaling have uncovered key developmental processes; however, these approaches cause embryo-wide FGF signaling perturbations, rendering assessment of cell-autonomous versus non-autonomous requirements for FGF signaling in individual processes difficult. Here, we created the novel transgenic line *fgfr1*-*dn*-*cargo*, encoding dominant-negative Fgfr1 with fluorescent tag under combined *Cre/lox* and heatshock control to provide spatio-temporal perturbation of FGF signaling. Validating efficient perturbation of FGF signaling by *fgfr1*-*dn*-*cargo* primed with ubiquitous CreERT2, we established that primed, heatshock-induced *fgfr1*-*dn*-*cargo* behaves akin to pulsed treatment with the FGFR inhibitor SU5402. Priming *fgfr1*-*dn*-*cargo* with CreERT2 in the lateral plate mesoderm, we observed selective cardiac and pectoral fin phenotypes without drastic impact on overall embryo patterning. Harnessing lateral plate mesoderm-specific FGF inhibition, we recapitulated the cell-autonomous and temporal requirement for FGF signaling in pectoral fin outgrow, as previously hypothesized from pan-embryonic FGF inhibition. Altogether, our results establish *fgfr1*-*dn*-*cargo* as a genetic tool to define the spatio-temporal requirements for FGF signaling in zebrafish.

## Introduction

Fundamental steps during vertebrate development are controlled by an evolutionary ancient family of secreted, diffusible polypeptides termed fibroblast growth factors (FGFs). The FGF family in mammals comprises 22 members (Huang and Stern, 2005), while in teleosts the number of FGF genes has expanded even further to 27 homologs/paralogs (Bökel and Brand, 2013). FGF ligands interact with cell surface-located FGF receptor tyrosine kinases (RTKs) on signal-receiving cells (Ornitz and Itoh, 2001; Plotnikov et al., 2000). In vertebrates, four FGF receptor (FGFR) genes (*Fgfr1*-*4*) have been identified(Itoh and Konishi, 2007). FGF ligand binding induces receptor dimerization, followed by tyrosine kinase activation by trans-phosphorylation, and primarily activation of the Ras/MAPK, PLC/Ca^2+^, and PI3 kinase/Akt cascades (Böttcher and Niehrs, 2005). The vast number of individual ligands with complex spatio-temporal expression patterns and the complex downstream effects of FGF signaling have greatly complicated the study of this key signaling pathway in vertebrate development. The protein tyrosine kinase inhibitor compound SU5402 prevents trans-phosphorylation by competing with ATP at the FGFR’s catalytic domain, allowing to experimentally inhibit FGF signaling (Mohammadi et al., 1997). Embryo-wide FGF perturbation during gastrulation results in aborted development of mesodermal and posterior structures, while pathway over-activation causes embryo dorsalization (Deng et al., 1994; Kroll and Amaya, 1996; Oki et al., 2010; Ota et al., 2009; Sun et al., 1999). Additionally, functional redundancy of FGFs and FGFRs complicate the evaluation of the impact of this signaling pathway on developmental processes (Ornitz and Itoh, 2015). The current challenge remains to separate the cell-autonomous activities of molecular pathways from their roles in global embryo patterning.

To date, FGF signaling studies focusing on the development of individual tissues have used traditional genetic perturbation methods in zebrafish including morpholinos, chemical genetics, and mutants, resulting in FGF signaling perturbations in the whole embryo. For example, the mutant *acerebellar (ace/fgf8a)* features perturbed heart formation, revealed by reduced cardiac marker expression and aberrant chamber development (Marques et al., 2008; Reifers et al., 2000), yet also displays axis formation defects and lacks the midbrain-hindbrain boundary and the cerebellum (Brand et al., 1996; Reifers et al., 1998). Correspondingly, FGF signaling inhibition by SU5402 treatment does not only effect the early cardiac gene expression induction, establishment of the myocardial progenitor pool, and organ territories within the ALPM, but also leads to shortening of posterior axis structures and defects in brain patterning (Reifers et al., 2000; Simões et al., 2011).

In zebrafish, FGF signaling has been implicated in directing cardiac versus either endothelial/myeloid or forelimb cell fates (Marques et al., 2008; Simoes et al., 2011). Surprisingly, in contrast to Fgf8 function in the mouse and chick limb buds, *ace* mutants do not feature any pectoral fin defects (Crossley et al., 1996; Lewandoski et al., 2000; Moon and Capecchi, 2000; Reifers et al., 1998). Instead Fgf24, Fgf10, and Fgf16 have been implicated in zebrafish pectoral fin development, where they act during mesenchyme compaction occurring between 18-28 hpf and on AER establishment after 36 hpf (Fischer et al., 2003; Nomura et al., 2006; Norton et al., 2005), but not during early limb induction (Mercader et al., 2006). While FGF signaling significantly contributes to cardiac and limb development, more precise spatio-temporally controlled perturbation of FGF signaling by genetic means would provide a crucial approach to dissect the cell-autonomous functions and windows of action for FGF signaling.

Previous work has established genetically controlled signaling inhibitors based on transgenes to modulate FGF signaling. Mutant FGFR with a non-functional cytoplasmic kinase domain can act as dominant-negative signaling inhibitor by forming unproductive heterodimers with native FGFR molecules, leading to their sequestration (Amaya et al., 1991; Ledda and Paratcha, 2007; Ota et al., 2009; Ullrich and Schlessinger, 1990). In zebrafish, dominant-negative FGFR1 efficiently inhibits FGF signaling when ubiquitously driven by a heatshock-controlled *hsp70:dn*-*fgfr1* transgene (Lee et al., 2005). Beyond temporal control of signaling perturbation akin to timed addition of chemical inhibitors, spatial control of such transgene-encoded signaling modulators remains a key challenge. In addition to the directly acting Gal4/*UAS* system, the Cre/*lox* system provides true uncoupling of transgene expression for precise spatio-temporal control: a Cre-expressing transgene triggers excision of a *lox*-flanked (floxed) cassette, resulting in productive expression of a signaling modulator as the transgenic cargo that becomes uncoupled from Cre activity (Branda and Dymecki, 2004; Rossant and Nagy, 1995). This principle has been harnessed in a limited number of studies to drive transcription factors, chromatin modulators, and signaling effectors (Carney and Mosimann, 2018).

In addition to achieving tight transgene control without any potentially detrimental leaky expression, the kinetics of transgene-based signaling modulators need to be fast enough to reach functional levels in the embryo to elicit an inhibitory response. Transgenic drivers based on ubiquitously active promoter elements such as *beta*-*actin* or *ubiquitinB* enable broad transgene expression, yet transgene expression upon Cre-mediated activation requires hours or even days to reach detectable levels (Carney and Mosimann, 2018; Chen et al., 2017; Hans et al., 2009; Mosimann et al., 2011). Heatshock-triggered transgenes have more rapid kinetics, and transgenic cargo under *hsp70l* promoter control is detectable less than one hour after heatshock (Hans et al., 2011; Hesselson et al., 2009).

Here, we generated and functionally evaluated a novel transgenic zebrafish line carrying a fluorescently marked, Cre/*lox*- and heatshock-controlled transgene based on the dominant-negative form of FGF receptor 1a (FGFR1a) to spatio-temporally perturb FGF signaling. The resulting transgene *Tg(*-*1.5hsp70l:loxP*-*STOP*-*loxP*-*fgfr1a*-*dn*-*2A*-*Cerulean*-*CAAX)*, abbreviated as *fgfr1*-*dn*-*cargo*, enables cell type-specific priming by CreERT2 recombinase and subsequent heatshock-controlled expression of FGFR1a dominant-negative protein with concomitant blue membrane labeling. We establish that *fgfr1*-*dn*-*cargo* triggered under its controlling stimuli results in pulsed inhibition of FGF target gene control akin to pulsed SU5402-mediated chemical inhibition of the pathway. When applied to perturb FGF signaling during cardiac and pectoral fin formation in the LPM, *fgfr1*-*dn*-*cargo* triggered in the descendants of *drl*-expressing LPM cells resulted in selective heart and pectoral fin defects without other pan-embryonic FGF loss-of-function phenotypes. Combining this spatio-temporal inhibition, we established two windows of LPM-autonomous FGF sensitivity for pectoral fin formation. Taken together, *fgfr1*-*dn*-*cargo* provides a versatile transgenic for spatio-temporal inhibition of FGF signaling activity applicable to broad developmental and regenerative contexts in zebrafish.

## Results

### A floxed and heatshock-dependent transgene to drive dominant-negative FGFR1a in zebrafish

Hetero-dimerization of FGFR with constitutive-active or dominant-negative forms of FGFR can sequester the native receptors and consequently modulate signaling activity, as achieved in ubiquitously active transgenes (Lee et al., 2005). To achieve spatio-temporal control over FGF signaling inhibition, we incorporated a dominant-negative Fgfr1a (Fgfr1a-dn) version carrying an inactivating mutation in its kinase domain (Lee et al., 2005; Ota et al., 2010) and a fluorescent marker coupled in a Cre/*lox*- and heatshock-controllable *Tol2* transgene, akin to the HOTcre approach (Hesselson et al., 2009). The resulting transgene *Tg(*-*1.5hsp70l:loxP*-*STOP*-*loxP*-*fgfr1a*-*dn*-*2A*-*Cerulean*-*CAAX*, *α*-*crystallin:YFP*) (Figure 1A) i) primes *fgfr1a*-*dn* expression upon Cre-mediated *loxP* recombination in the cell type and at the time of choice; ii) subsequent heatshock treatment activates the transgene at the desired perturbation stage; and iii) membrane-bound blue-fluorescent Cerulean-CAAX marks all cells with successfully activated transgene. Due to the strong position sensitivity of *lox* cassette transgenes (Felker and Mosimann, 2016; Mosimann and Zon, 2011; Mosimann et al., 2011), we screened over a dozen transgenic insertions before establishing one stable transgenic line *Tg(*-*1.5hsp70l:loxP*-*STOP*-*loxP*-*fgfr1a*-*dn*-*2A*-*Cerulean*-*CAAX^VII^)*, from here-on called *fgfr1*-*dn*-*cargo*, that showed reproducible recombination and transgene expression efficiency.

**Figure 1:**
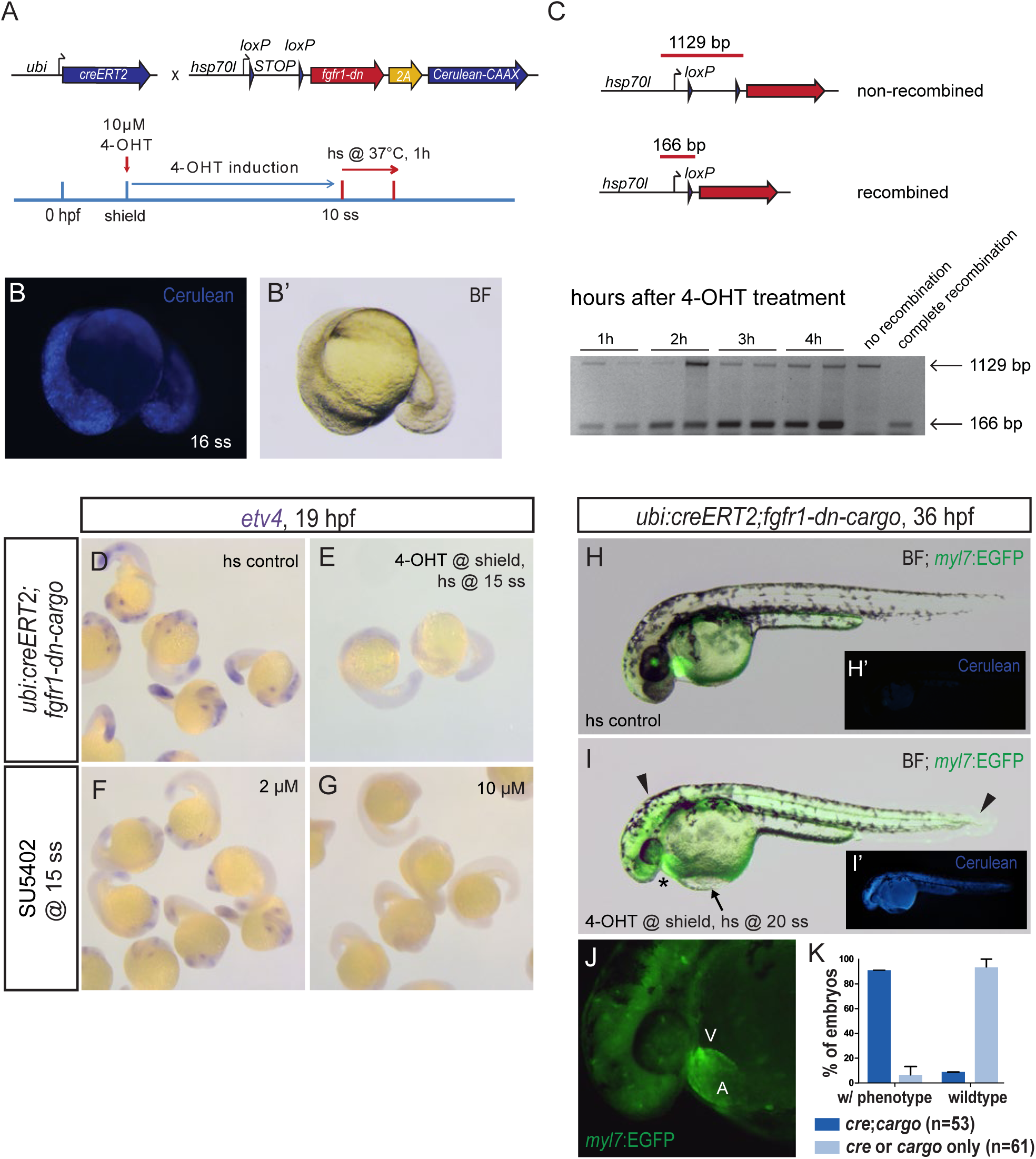
Global FGF signaling perturbation using the *fgfr1*-*dn*-*cargo* transgenic line. (**A**) Schematic showing the crosses and a representative treatment scheme for ubiquitous genetic FGF signaling perturbation using the *fgfr1*-*dn*-*cargo* transgenic line. Note that *fgfr1*-*dn*-*cargo* contains a *2A*-linked *Cerulean*-*CAAX* ORF. (**B,B’**) Ubiquitous *fgfr1*-*dn*-*cargo* activation in 4-OHT- and heatshock-treated *ubi:creERT2*;*fgfr1*-*dn*-*cargo* transgenic embryos as shown in schematic (**A**) leads to ubiquitous, but mosaic Cerulean-CAAX (Cerulean) expression. (**C**) CreERT2-mediated ubiquitous excision of the *loxP*-flanked *STOP* cassette in *ubi:creERT2*;*fgfr1*-*dn*-*cargo* transgenics 4-OHT-induced during early somitogenesis as detected by PCR (166 bp after recombination, 1129 bp when un-recombined). Excision of the *STOP* cassette occurs within one hour and gradually increases up to four hours after 4-OHT treatment. (**D-G**) Heatshock (hs) controls, *ubi:creERT2*;*fgfr1*-*dn*-*cargo* embryos induced with 4-OHT at shield stage and heat-shocked at 15 ss, and wildtype embryos treated with 2 or 10 μM SU5402 at 15 ss were fixed at 19 hpf and stained for *etv4* expression via mRNA *in situ* hybridization. Ubiquitous *fgfr1*-*dn*-*cargo* activation and concentration-dependent SU5402 treatment abolish *etv4* expression as a read-out for FGF signaling activity. (**H,I**) Overlay of EGFP expression on a brightfield (BF) image of a 36 hpf heatshock control and *ubi:creERT2*;*fgfr1*-*dn*-*cargo* embryo induced with 4-OHT at shield stage and heatshock-treated at 20 ss. Embryos with ubiquitous expression of Fgfr1-dn during late somitogenesis have a miss-looped heart (asterisk) as well as head defects and malformations in posterior tail structures (arrowheads). The heart malformations also manifest in blood pooling in front of the inflow tract of the heart, leading to a visible edema on top of the yolk (arrow). (**H’,I’**) Cerulean-CAAX expression in control and signaling-perturbed embryos (**H,I**). (**J**) 7x magnification of the heart of a 4-OHT-induced and heatshock-treated *ubi:creERT2*;*fgfr1*-*dn*-*cargo* double transgenic (**I**) showing heart defects with a large atrium (A) and diminished ventricle (V); green background fluorescence is caused by bleed-through of Cerulean fluorescence (see also **I’**). (**K**) Quantification of phenotypes resulting from global FGF signaling perturbation in genetically perturbed double-transgenic *ubi:creERT;fgfr*-*dn*-*cargo* embryos and single-transgenic heatshock controls treated as indicated in (**I**).

To test the general functionality of the stable *fgfr1*-*dn*-*cargo* line, we first ubiquitously primed *fgfr1*-*dn* expression through recombination using the *ubi:creERT2* driver (Mosimann et al., 2011): treating embryos with 4-OHT at shield stage and inducing transcriptional activation of the *fgfr1*-*dn*-*cargo* cassette with heatshock treatment at 10 somite stage (ss) resulted in ubiquitous mosaic Cerulean-CAAX expression in double-transgenic embryos, indicating successful transgene expression (Figure 1A, 1B). To analyze the precise dynamics of 4-OHT-mediated recombination of the *fgfr1*-*dn*-*cargo* transgene, we analyzed excision of the *loxP*-flanked *STOP* cassette from genomic DNA via PCR. Following ubiquitous CreERT2 recombinase activity, we robustly detected successful recombination as soon as 1 hour after 4-OHT treatment and more efficient recombination 2 hours after 4-OHT treatment, indicating fast *in vivo loxP* recombination in line with previous reports (Hans et al., 2009) (Figure 1C).

We next tested the efficacy of FGF signaling perturbation by comparing expression of the direct FGF downstream target *etv4/pea3* (Raible and Brand, 2001; Roehl and Nüsslein-Volhard, 2001) upon global *fgfr1-dn-cargo* transgene activation or chemical inhibition of endogenous FGFRs with the established compound SU5402 (Mohammadi et al., 1997). In Cerulean-CAAX-expressing *ubi:creERT2;fgfr1*-*dn*-*cargo* double-transgenics that we had primed with 4-OHT at shield stage (6 hpf) and heatshock-treated at 15 ss (16.5 hpf), we observed by mRNA *in situ* hybridization a complete loss of *etv4* expression at 20-25 somite stage (19 hpf) (Figure 1D,E); we saw an equivalent effect on *etv4* expression in wildtype embryos of the same stage when treated at 15 ss with high concentrations of SU5402 (Figure 1F,G). Activating ubiquitous Fgfr1-dn expression at 20 ss by heatshock treatment, we further detected morphological defects corresponding to phenotypes previously described upon global FGF signaling perturbations during late somitogenesis (Marques et al., 2008): a miss-looped heart with large atria and a diminished ventricle, lack of blood flow, plus head and posterior tail malformations (Figure 1H-K, n=53). Altogether, these results indicate that our *fgfr1-dn-cargo* line provides a functional zebrafish transgene for perturbing the FGF signaling pathway by driving dominant-negative Fgfr1a.

### Dynamics of FGF signaling perturbation from *fgfr1*-*dn*-*cargo* resembles pulsed SU5402 treatment

Heatshock-mediated transgene induction generates a pulse of *hsp70l* promoter-driven transcription, resulting in transient expression of the controlled transgene. We therefore hypothesized that our *fgfr1*-*dn*-*cargo* transgene provides pulsed inhibition of FGF signaling. To analyze the temporal dynamics of transgene expression, we compared the impact on *etv4* expression in i) 4-OHT- and heatshock-treated *ubi:creERT;fgfr1*-*dn*-*cargo* double-transgenics; ii) embryos exposed to SU5402 for a four-hour pulse before washing out the drug; and iii) Cerulean negative single-transgenic controls (Figure 2A). We chose 10-11 ss (approx.14-15 hpf) to initiate FGF inhibition as *etv4* expression is then easily detectable by mRNA *in situ* hybridization in several regions of the developing embryo (Figure 2B-F).

**Figure 2:**
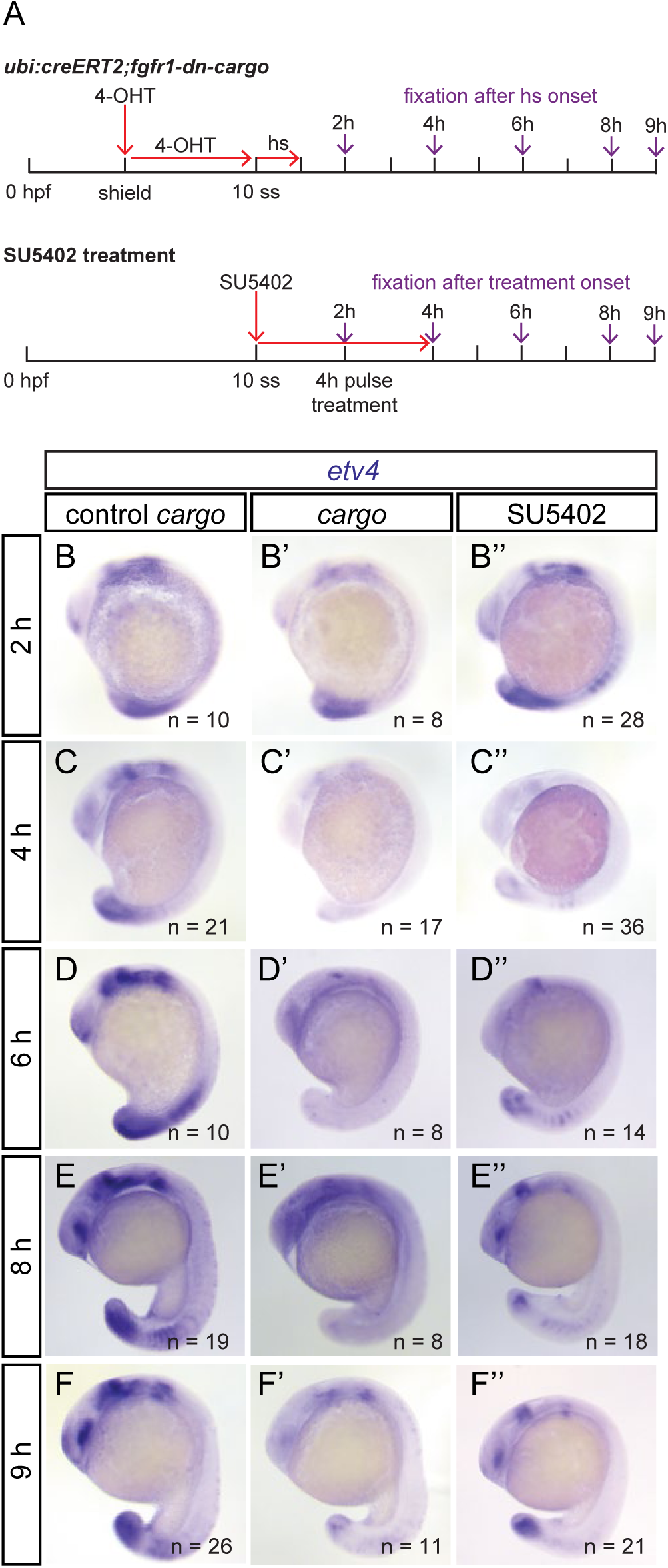
Temporal dynamics of genetic FGF signaling perturbation using *fgfr1*-*dn*-*cargo*. (**A**) Schematic showing the timeline of treatments for genetic and chemical FGF signaling perturbations and timepoints of embryo fixation for signaling activity read-outs. Red arrows indicate timepoints and duration of drug (10 μM 4-OHT and 5 μM SU5402) and heatshock treatments, violet arrows mark timepoints of embryo fixation after treatment onset (heatshock or SU5402 application). (**B-F**) Representative embryos stained for *etv4* mRNA expression via *in situ* hybridization in Cerulean negative single-transgenic controls (control *cargo*), *ubi:creERT2;fgfr1*-*dn*-*cargo* (*cargo*), and SU5402-treated embryos; lateral views, anterior to the top. Numbers “n” indicate individual embryos stained and analyzed for each condition. (**B,C**) After FGF inhibition was initiated at mid-somitogenesis (10 ss), *etv4* expression in *ubi:creERT2;fgfr1*-*dn*-*cargo* and SU5402-treated embryos was decreased two hours and absent four hours after treatment onset. (**D-F**) *etv4* expression started recovering two hours after SU5402-treatment was stopped (six hours after initiation of a four-hour pulse) and eight hours after heatshock in *ubi:creERT2;fgfr1*-*dn*-*cargo* double-transgenics. Nine hours after treatment onset, *etv4* expression recovered to a large extent in genetically and pulse-treated chemically perturbed embryos (**F**).

Both in *ubi:creERT2;fgfr1*-*dn*-*cargo* double-transgenics and in SU5402-treated embryos, strong reduction of *etv4* expression became detectable within two hours after treatment, and was completely absent four hours after treatment (Figure 2B-C). *etv4* expression remained broadly absent up to six hours after heatshock treatment or SU5402 addition (two hours after wash-out) (Figure 2D). Eight and nine hours following transgene activation or SU5402 treatment (corresponding to four and five hours after washout), respectively, *etv4* expression was still notably reduced, but expression started to recover in both conditions, with possibly slightly slower recovery in *ubi:creERT2;fgfr1*-*dn*-*cargo* embryos (Figure 2E,F). The dynamics of *etv4* expression in *ubi:creERT2;fgfr1*-*dn*-*cargo* double-transgenics and SU5402 pulse-treated embryos reveal that ubiquitous *fgfr1*-*dn*-*cargo* transgene activation resembles chemical FGFR inhibition in respect to strength and onset dynamics. Further, the upregulation of *etv4* expression within eight hours after transgene activation is consistent with the notion that FGF signaling perturbation in the *fgfr1*-*dn*-*cargo* line does not occur indefinitely but as a pulse.

We further sought to analyze FGF signaling activity in embryos that were genetically or chemically perturbed during gastrulation and initiated *fgfr1*-*dn*-*cargo* expression or SU5402 treatment at shield stage (Figure 3A). Under these conditions, we did not detect any embryos with complete absence of FGF signaling activity, as read out by *etv4* expression; nonetheless, we documented reduced FGF signaling three and, more prominently, four hours after either genetic or chemical FGF signaling perturbation (Figure 3B,C). Consequently, experiments aiming for FGF signaling perturbations during gastrulation ought to consider slower dynamics and milder effects on FGF signaling attenuation when using the *fgfr1*-*dn*-*cargo* transgene.

**Figure 3:**
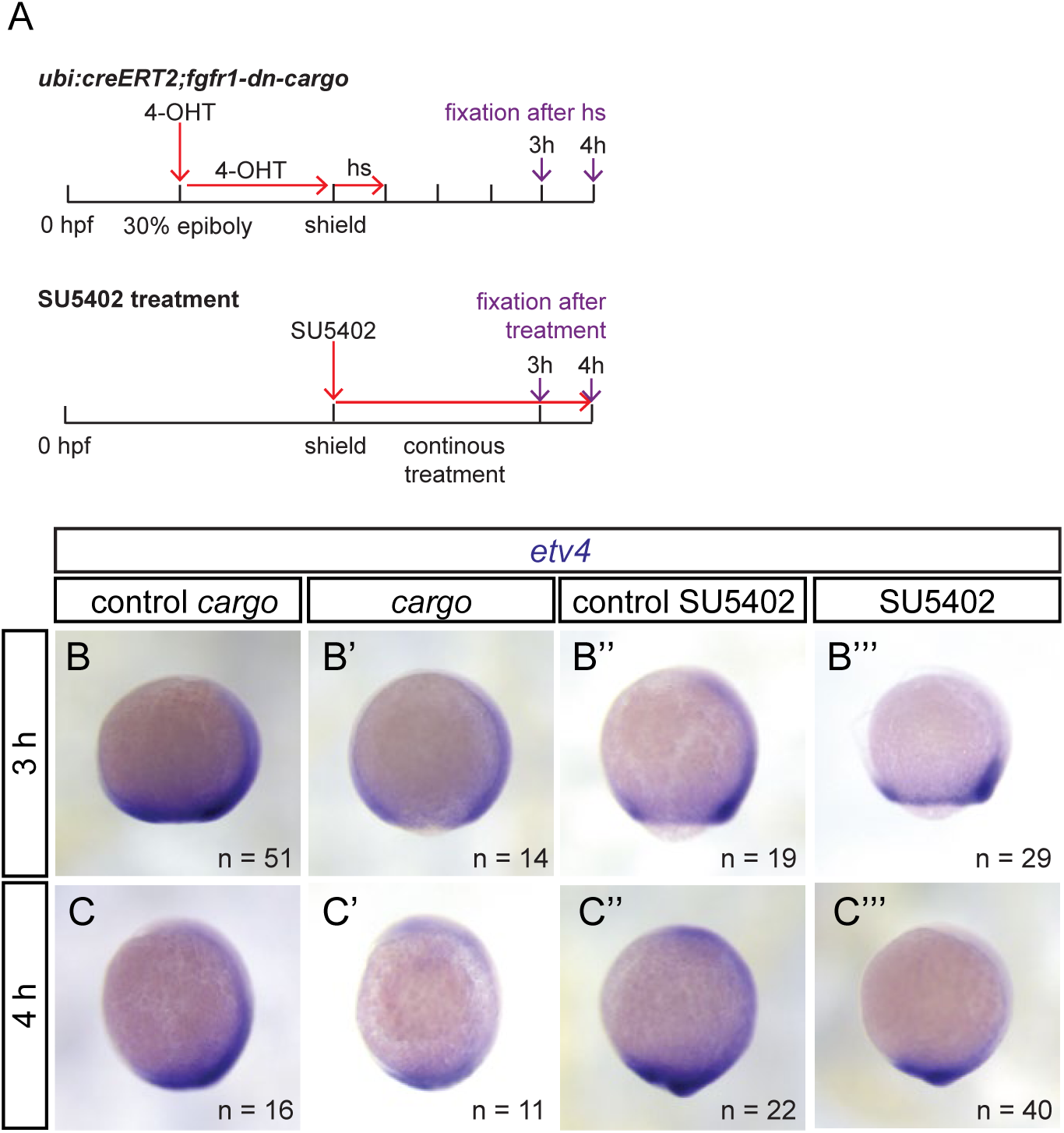
Dynamics of genetic and chemical FGF signaling perturbation during gastrulation. (**A**) Schematic showing the timeline of treatments for genetic and chemical FGF signaling perturbations and timepoints of embryo fixation for signaling activity read-outs. Red arrows indicate timepoints and duration of drug (10 μM 4-OHT and 5 μM SU5402) and heatshock treatments, violet arrows mark timepoints of embryo fixation after treatment onset (heatshock or SU5402 application). Note that Cerulean-CAAX expression could not be observed between one to two hours post-heatshock, thus *etv4* expression analysis is only shown after three hours. (**B,C**) Representative embryos stained for *etv4* mRNA expression with *in situ* hybridization in Cerulean negative single-transgenic controls (control cargo), Cerulean-positive *ubi:creERT2;fgfr1*-*dn*-*cargo* double-transgenics (*cargo*), untreated wildtype controls (control SU5402), and SU5402-treated embryos; lateral views, anterior to the top. “n” indicates individual embryos stained and analyzed for each condition. *etv4* expression in *ubi:creERT2;fgfr1*-*dn*-*cargo* and SU5402-treated embryos was decreased three and four hours after treatment at shield stage, but never completely lost as at later stages (see also **Figure 2 C,D**).

### Perturbation of FGF signaling in restricted cell lineages

To perform spatio-temporally controlled FGF signaling inhibition using our *fgfr1*-*dn*-*cargo* transgene, we next crossed it to *drl:creERT2* transgenic zebrafish. *drl*-based reporters label the forming LPM from late gastrulation to early somitogenesis before confining expression to cardiovascular lineages (Henninger et al., 2017; Hess et al., 2018; Mosimann et al., 2015). To test if *drl:creERT2*-mediated recombination of the *fgfr1*-*dn*-*cargo* transgene sufficiently triggers heatshock-dependent Fgfr1-dn expression in the developing LPM, we performed *in toto* SPIM-imaging of Cerulean-CAAX as a proxy for transgene expression in *drl:creERT2;fgfr1*-*dn*-*cargo* double-transgenics. After 4-OHT-induction at 30% epiboly and heatshock treatment during somitogenesis (10 ss) we observed LPM-confined mosaic Cerulean expression in the entire LPM at 14 ss (Figure 4A-C).

**Figure 4:**
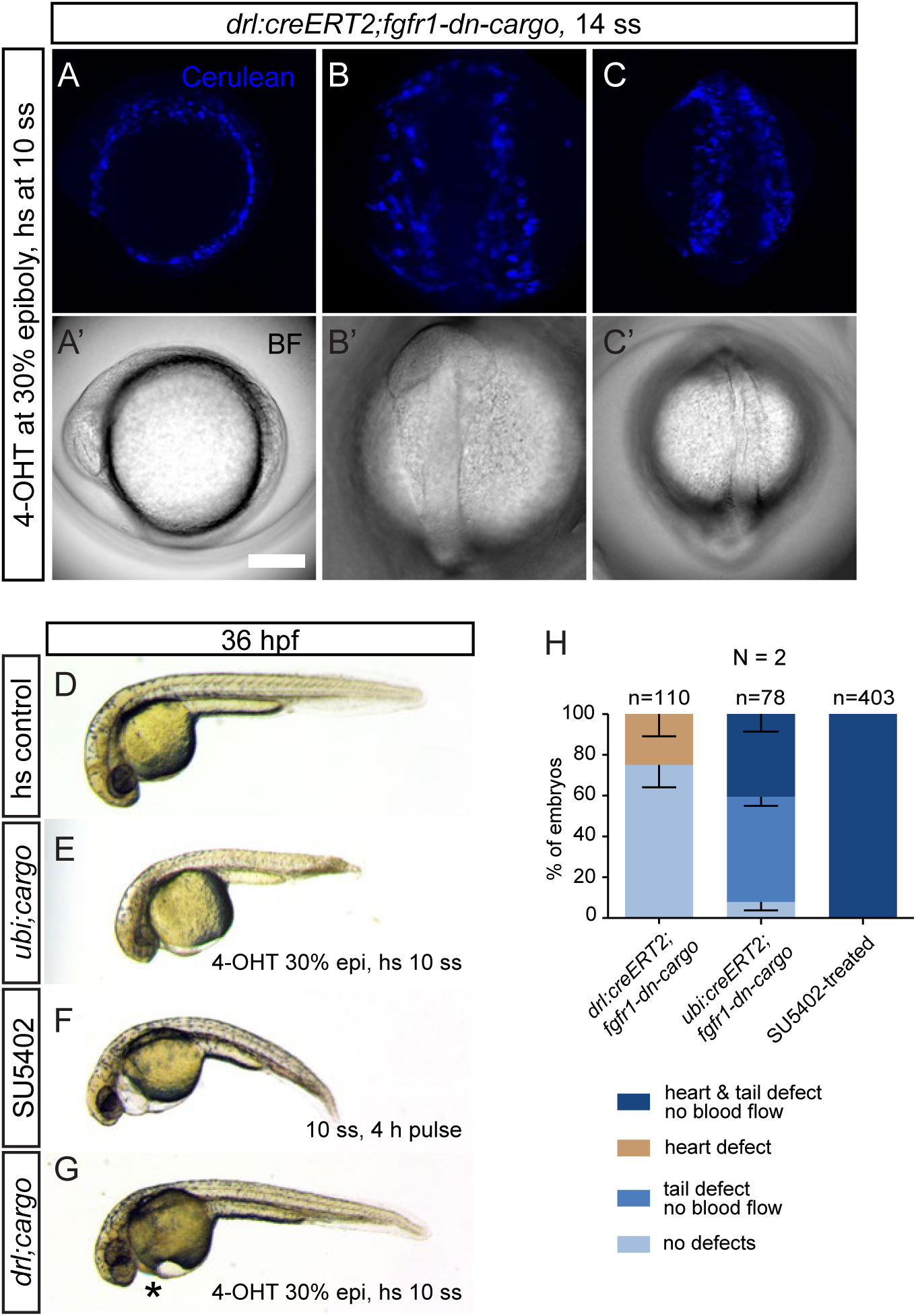
Tissue-specific FGF signaling perturbation in the developing LPM. (**A-C**) Maximum intensity projection and bright field images of an *in toto* SPIM-imaged *drl:creERT2;fgfr1*-*dn*-*cargo* double-transgenic embryo. The cell membrane of all Fgfr1-dn-expressing cells is fluorescently labeled with Cerulean-CAAX after heatshock treatment revealing mosaic transgene expression throughout the developing LPM. (**A**) lateral view, anterior to the top; (**B**) dorsal view of the anterior embryo, anterior to the top; (**C**) dorsal view of the posterior embryo, anterior to the top. (**D-G**) Brightfield images of globally or tissue-specifically FGF signaling perturbed embryos, lateral views, anterior to the left. (**E,F**) Ubiquitously Fgfr1-dn-expressing or SU5402-treated embryos show severe cardiovascular defects accompanied by body axis shortening and defects in posterior tail formation. (**G**) Selective cardiovascular phenotypes (asterisk) with no apparent body axis deformations are apparent after tissue-specific FGF signaling perturbation in *drl* expressing descendants. (**H**) Quantifications of phenotypes observed after *drl:creERT2;fgfr1*-*dn*-*cargo*, *ubi:creERT2;fgfr1*-*dn*-*cargo*, or SU5402-mediated FGF signaling inhibition, n indicates the number of individual embryos analyzed per condition, N indicates the number of individual experiments performed.

Next, we compared phenotypes of SU5402-perturbed wildtype embryos to double-transgenic embryos for *ubi:creERT2;fgfr1*-*dn*-*cargo* (ubiquitous FGF perturbation) and *drl:creERT2;fgfr1*-*dn*-*cargo* (LPM-specific FGF perturbation), respectively. We primed ubiquitous and lineage-specific *loxP* recombination with 4-OHT at 30% epiboly to shield stage (to target the earliest progenitors expressing *drl:creERT2*), activated *fgfr1*-*dn*-*cargo* expression via heatshock at 10-11 ss, and performed phenotype observations at 36 hpf. Defects in embryos following i) ubiquitous *fgfr1*-*dn*-*cargo* transgene activation (n=78), or ii) a 4-hour pulse of SU5402 at the same stage (n=403) resembled the phenotypes seen before (Figure 1I): a miss-looped heart, lack of blood flow, as well as head and tail defects (Figure 4D-F,H). In contrast, in embryos perturbed selectively in the *drl* descendants, we observed milder phenotypes mainly characterized by heart defects apparent through blood pooling and edema at the cardiac cavity (n=24) or no phenotypes (n=86) (Figure 4G,H, n=110, N=2). Although we also detected defects in the posterior endothelium, we never observed a complete block of blood circulation nor posterior tail defects in *drl:creERT2;fgfr1*-*dn*-*cargo* double-transgenics.

Expression of a dominant-negative receptor could potentially act non-cell-autonomously by scavenging FGF ligand from the extracellular space, rendering it unavailable to neighboring cells. We therefore revisited the expression of *etv4*: in *drl:creERT2;fgfr1*-*dn*-*cargo* double-transgenics, 4-OHT-treated at shield stage and heatshock-treated at 10-11 ss, we did not detect any overt changes to *etv4* expression up to five hours after transgene activation (Supplementary Figure 1). Observing identically treated embryos at 36 hpf (more than 21.5 hours post-heatshock), expression of the FGF targets *etv4*, *spry4*, and *dusp6* remained broadly normal, in contrast to the effect of constant exposure to SU5402 (Figure 5, Supplementary Figures 2,3). Nonetheless, we observed reduced expression of *etv4* and *spry4* in the pectoral fin buds of *drl:creERT2;fgfr1*-*dn*-*cargo* double-transgenic embryos (Figure 5 A,C,F, Supplementary Figure 2), while *dusp6* expression remained seemingly wildtype (Supplementary Figure 3). We further detected these *etv4* and *spry4* phenotypes upon similarly timed ubiquitous FGF inhibition in *ubi:creERT2;fgfr1*-*dn*-*cargo* double-transgenics (n=50/60) and upon pulsed SU5402 treatment (n=21/56), yet both these conditions also perturbed non-LPM domains of the probed FGF targets (Figure 5 D-F, Supplementary Figures 2, 3). Continuous treatment with SU5402 caused complete loss of the pectoral fin domain shown by *etv4*, *spry4*, and *dusp6* and also the anticipated overall loss of their expression concomitant with perturbed embryo morphology (Figure 5 E,F, Supplementary Figures 2, 3).

**Figure 5:**
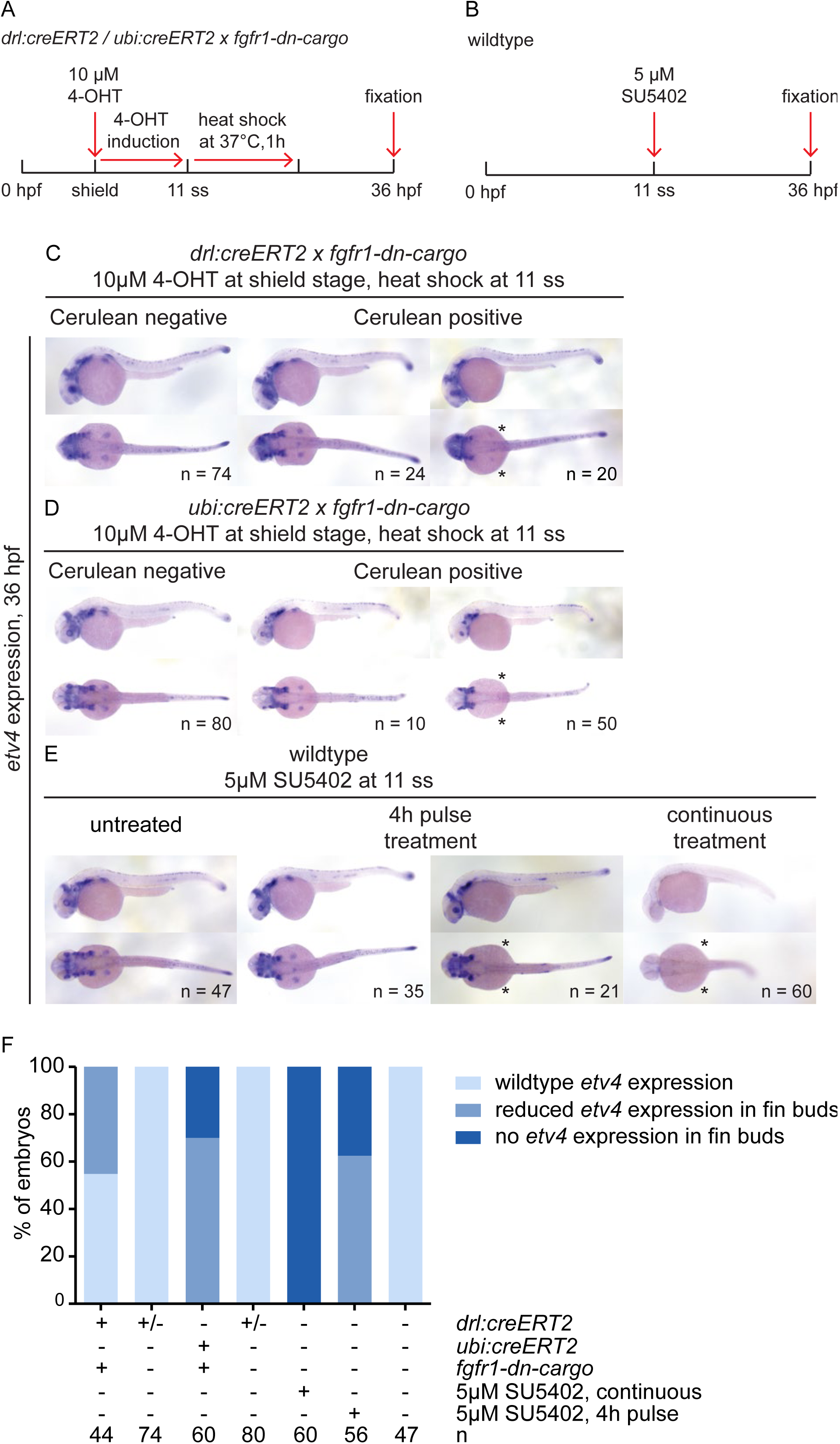
*etv4* expression in tissue-specifically and globally FGF signaling-perturbed embryos at 36 hpf. (**A, B**) Transgenic embryos were treated with 4-OHT at shield stage and heatshock-treated at 11 ss (**A**), while wildtype embryos were treated with SU5402 either continuously or for a four-hour pulse for comparison; (**B**) expression of the FGF target gene etv4 assayed at 36 hpf. (**C-E**) *etv4* expression in the different conditions. *etv4* expression was grossly unaffected by LPM-specific perturbation using *drl:creERT2* priming *fgfr1*-*dn*-*cargo* (**C**, compare Cerulean-negative to Cerulean-positive, *fgfr1*-*dn*-expressing embryos), with notable exception of pectoral fin expression that was absent in a cohort of *fgfr1*-*dn*-expressing embryos (asterisks). *etv4* was also grossly unaffected after ubiquitous FGF perturbation in cohorts of *ubi:creERT2;fgfr1*-*dn*-*cargo* embryos (**D**,) and SU5402 pulse-treated embryos (**E**), while pectoral fin expression was again affected (asterisks in **D**, **E**). Continuous FGF signaling inhibition with SU5402 as per indicated conditions (**B**) broadly inhibited *etv4* expression (asterisks in **E**). (**F**) Quantification of *etv4* expression in embryos subjected to embryo-wide or tissue-specific signaling perturbations; n indicates numbers of individual embryos analyzed for each condition.

These data indicate that tissue-specific Fgfr1-dn expression does not cause pan-embryonic attenuation of FGF signaling activity, arguing against a broad removal of FGF ligand from the extracellular space by the forced tissue-specific expression of dominant-negative Fgfr1. Our observations further suggest a lasting pectoral-specific effect of FGF perturbation for the window of activity starting at 10-11 ss, possibly influenced by the level of mosaicism following floxed STOP cassette excision.

### Temporally distinct requirements for FGF signaling control pectoral fin development in the LPM

While dispensable for initial fin bud induction (Mercader et al., 2006), perturbed FGF signaling starting from approximately 18 ss (18 hpf) and beyond interferes with proper pectoral fin formation, as analyzed in genetic mutants (Fischer et al., 2003; Nomura et al., 2006; Norton et al., 2005) and pan-embryo inhibition using SU5402 (Mercader et al., 2006; Prykhozhij and Neumann, 2008). *fgfr1*-*dn*-*cargo* expression in the LPM at 11 ss caused a selective impact on FGF target gene expression in the pectoral fin buds (Figure 5, Supplementary Figures 2,3). We therefore sought to apply *fgfr1*-*dn*-*cargo* to recapitulate if and when LPM lineage-specific perturbation of FGF is sufficient to cause detectable phenotypes in the pectoral fins.

We again crossed heterozygous *drl:creERT2* males to heterozygous *fgfr1*-*dn*-*cargo* females, induced in their offspring CreERT2-mediated recombination with 4-OHT at 30% epiboly to maximize LPM priming, performed heatshock treatment to activate *fgfr1*-*dn*-*cargo* at discrete time points, and sorted embryos by Cerulean expression as double-transgenic (approx. 25%) versus *drl:creERT2* or *fgfr1*-*dn*-*cargo* siblings; we also performed the same treatment in independent wildtype controls (Figure 6A). At 48-56 hpf, we observed heart phenotypes based on obvious looping defects and development of a cardiac edema as previous studies (Marques et al., 2008) (Figure 6B,C); we in detail scored fin defects based on uni- or bilateral malformation up to complete fin loss (Figure 6D-F).

**Figure 6:**
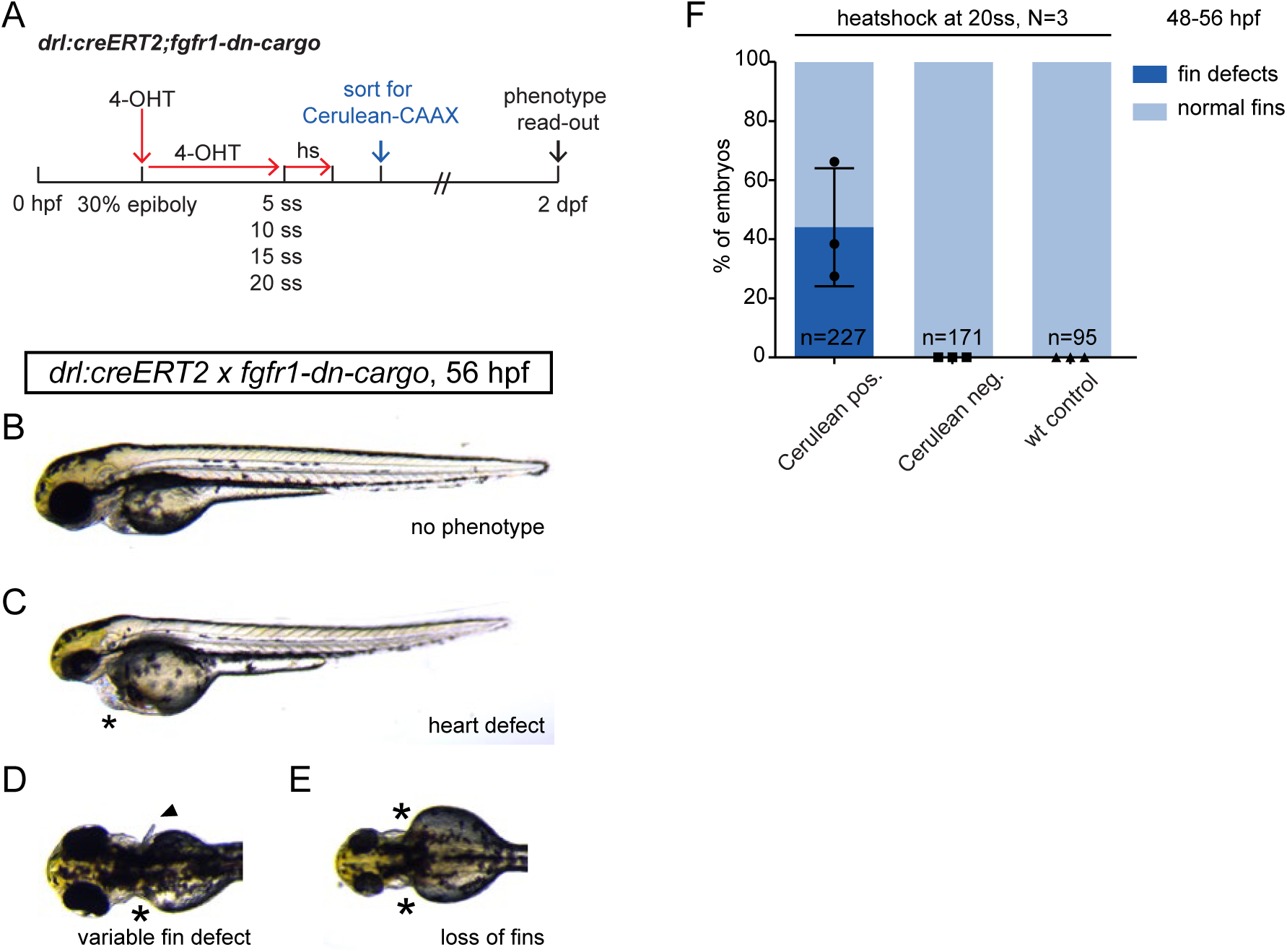
Temporal dissection of tissue-specific FGF signaling control of cardiac and fin development. (**A**) Schematic showing experimental timeline of tissue-specific FGF signaling perturbations at discrete developmental stages. 4-OHT induction at 30% epiboly induces Cre/*lox* recombination in earliest *drl*-expressing progenitors. Heatshock treatments at distinct timepoints throughout somitogenesis target different phases of cardiac and fin development. (**B-D**) Phenotypes scored in 2 dpf *drl:creERT2;fgfr1*-*dn*-*cargo* embryos treated as indicated in the schematic above. (**B,C**) Heart defects as scored through blood pooling and cardiac edema on top of the yolk (asterisks), lateral views, anterior to the top. (**D-E**) Fin defects were counted upon uni- or bilateral fin deformations (arrow head) and/or loss (asterisks). (**F-I**) Quantifications of phenotypes in *drl:creERT2;fgfr1*-*dn*-*cargo* double-transgenics heatshock-treated at different timepoints during somitogenesis. (**F**) LPM-specific FGF perturbation triggered by heatshock at 20 ss results in pectoral fin defects in on average 44.09% (s.d. 20.01, p=005, total n=227, three independent experiments) of double-transgenic embryos with prior Cerulean expression, while earlier heatshock timings caused no discernible fin defects. n indicates the number of individual embryos analyzed per condition, N indicates the number of individual experiments performed; statistics based on one-way Anova, multiple comparison, Tukey’s post test.

With LPM-restricted *fgfr1*-*dn*-*cargo* activation at 5 ss, 10 ss, 15 ss, and 20 ss, we consistently observed variable, yet reproducible cardiac edema in embryos after heatshock, (ranging from 11% to 26%), suggesting functional FGF perturbation consistent with previous observations (Marques et al., 2008). In contrast, we observed no visible pectoral fin defects in embryos with LPM-restricted *fgfr1*-*dn*-*cargo* activation at 5 ss, 10 ss, or 15 ss (n=58, 190, and 172, respectively, from three independent experiments each). Instead, while not fully penetrant, heatshock treatment at 20 ss resulted in uni- or bilaterally lost fins (44% of Cerulean-positive embryos, n=227, three independent experiments; Figure 6F). This observation upon transient, LPM lineage-focused FGF inhibition starting at 20 ss (approximately 19 hpf) coincides with previously reported mutant and chemical perturbation experiments that assigned the early critical window of FGF signaling during pectoral fin formation between 18-28 hpf (Fischer et al., 2003; Norton et al., 2005).

## Discussion

The precise spatio-temporal modulation of signaling pathways *in vivo*, most-desirably within selected cell lineages at developmental times of interest, remains challenging. Key technical issues are the strict control and kinetics of signaling-modulating transgene expression. To date, existing studies on FGF signaling in zebrafish have predominantly reached their conclusions using elegant mutant genetics and chemical whole-embryo perturbations, such as in the analysis of heart and pectoral fin formation (de Pater et al., 2009; Dong et al., 2007; Fischer et al., 2003; Marques et al., 2008; Mercader et al., 2006; Norton et al., 2005; Reifers et al., 2000). Nonetheless, the cell- or lineage-autonomous contribution of FGF signaling in these processes remains inferred. We have generated *fgfr1*-*dn*-*cargo*, a novel zebrafish line carrying a Cre/*lox*-controlled transgene driving dominant-negative Fgfr1a (Lee et al., 2005; Ota et al., 2009) expression to spatio-temporally block FGF signaling. Building on previous direct *hsp70l*-driven and Cre/lox-controlled approaches (Hans et al., 2011; Hesselson et al., 2009; Lee et al., 2005), control on two levels defines the cell lineage and time of FGF signaling perturbation: i) time of 4-OHT induction primes the transgene in the exact lineage as per used CreERT2 driver; ii) time of heatshock treatment determines the developmental stage affected by rapid, pulsed FGF signaling perturbation. The *2A*-*Cerulean*-*CAAX* cassette provides a read-out for successful transgene expression and assessment of mosaicism by fluorescent membrane labeling (Figure 1A,B).

Ubiquitous *fgfr1*-*dn*-*cargo* transgene activation results in phenotypes and changed expression of FGF target gene resembling global chemical FGFR inhibition with SU5402 in our hands (Figure 1 H-K, Figure 4 E,F) and as previously reported (Brand et al. 1996, Marques et al, 2008, Reifers et al., 2000). Recombination in our *fgfr1*-*dn*-*cargo* line enables fast experimental timelines, permitting transgene activation one to two hours post 4-OHT treatments (Figure 1 C). Moreover, also akin to administration of SU5402, we document successful FGF perturbation within short timeframes after heatshock-induced transgene activation (Figure 1 D-G, Figure 2B,C). These results establish the *fgfr1*-*dn*-*cargo* transgenic as viable genetic tool to genetically block FGF signaling in discrete developmental timepoints and cell types.

While highly potent post-gastrulation, we also documented only partial inhibition of FGF signaling activity during early gastrulation stages using *fgfr1*-*dn*-*cargo*, yet also following SU5402-mediated chemical perturbation (Figure 3). Due to the maternal contribution of CreERT2 driven by the *ubi* promoter (Mosimann et al., 2011) and since chemical perturbation yielded similar results, we consider low levels of *loxP* recombination an unlikely cause for the incomplete inhibition. Instead, incomplete FGF signaling inhibition may be due to strong FGF signaling activity during gastrulation that is potentially already established through high levels of maternally contributed FGFRs and may not be easily overcome with our transgene. Thus, studies aiming to elucidate spatio-temporal requirements during gastrulation should consider different dynamics than presented for somitogenesis embryos above.

Expression of our *fgfr1*-*dn*-*cargo* transgene is under the control of the *hsp70l* promoter (Figure 1A) that has been successfully used for driving *loxP*-governed transgenes (Hans et al., 2011; Hesselson et al., 2009). As the activity of the *hsp70l* promoter ceases post-heatshock treatment, *fgfr1*-*dn* transcription ceases after the heatshock response has faded. This is consistent with the pulsed nature of the perturbation we have observed with our transgene, with complete FGF signaling inhibition up to six hours and perturbed signaling up to nine hours post-heatshock comparable to pulsed SU5402 treatments (Figure 2 D,E,F). Although *hsp70l* promoter transgenes can respond to other stimuli leading to non-conditional recombination at permissive temperatures (Hans et al., 2009 & 2010), we do not observe unspecific Cerulean expression in non-recombined heatshock controls (Figure 1H) or in recombined non-heatshock-treated transgenics (data not shown), revealing that the used *loxP* STOP cassette (Hesselson et al., 2009) is tight in our particular transgenic insertion. These observations are supported by the lack of phenotypes caused by perturbed FGF signaling in heatshock-only controls (Figure 4D). Nonetheless, *hsp70l* used to drive *fgfr1*-*dn*-*cargo* paired with suitable Cre/CreERT2 drivers provide a versatile combination for fast transgene activation to study fast-occurring FGF-dependent developmental processes. Moreover, temporally restricted inhibition by *fgfr1*-*dn*-*cargo* allows for detailed analysis of exact developmental windows with requirement for FGF signaling activity in a specific cell type.

FGFs expressed in the apical ectodermal ridge (AER) of the developing limb are essential to maintain a progenitor pool that ensures proximal-distal outgrowth and patterning of the mouse and chick limb from the LPM (Crossley et al., 1996; Fallon et al., 1994; Lewandoski et al., 2000; Moon and Capecchi, 2000; Niswander et al., 1993). In particular, ablation of Fgf8 in the AER lead to severe limb truncations while concurrent removal of Fgf8, 4, and 9 lead to a complete limbless phenotype, demonstrating redundancy and dose dependency (Mariani et al., 2008). Additionally, reciprocal AER FGF signaling activity induces Fgf10 expression in the distal limb mesenchyme which, in return, is necessary to maintain FGF signaling from the AER (Ohuchi et al., 1997; Sekine et al., 1999; Xu et al., 1998). In contrast, FGF signaling function in earlier steps of limb induction have been controversial. While application of Fgf8 to the chicken flank results in ectopic limb formation and Fgf8 expression in the intermediate mesoderm had been described to induce limb formation in chick, conditional removal of Fgf8 activity from the mouse intermediate mesoderm did not abrogate limb development (Crossley et al., 1996; Vogel et al., 1996). After priming in the developing LPM, triggering *fgfr1*-*dn*-*cargo* expression at 20 ss (19 hpf) resulted in disrupted pectoral fins (Figure 6D-F), while *fgfr1*-*dn*-*cargo* triggered at earlier time points caused no overt pectoral fin defects. Nonetheless, *fgfr1*-*dn*-*cargo* is functional in the LPM descendants, as i) expression of the FGF target genes *etv4* and *spry4* (Figure 5, Supplementary Figure 2), and less notably *dusp6* (Supplementary Figure 3), was reduced at 36 hpf in pectoral fin buds when *fgfr1*-*dn*-*cargo* was triggered at 10-11 ss (14-15 hpf), and ii) we observed cardiac phenotypes as reported for SU5402 treatment or genetic perturbations (de Pater et al., 2009; Marques et al., 2008; Pradhan et al., 2017; Reifers et al., 2000) (Figure 6C). Together, these observations support an LPM-autonomous requirement for FGF activity during the critical phase of pectoral fin bud outgrowth between 18-28 hpf in zebrafish, when FGF10 and FGF24 are active in the tissue (Fischer et al., 2003; Norton et al., 2005).

A caveat to interpreting Cre/*lox*-based phenotypes remains the variable mosaicism resulting from embryo-to-embryo variation in *loxP* cassette excision, as also observed in our work (Figure 4A-C). While our phenotypic observation focused on frequently observed overt changes to fin morphology, including complete absence of pectoral fins, we observed an incomplete phenotype penetrance (44% of all *drl:creERT2;fgfr1*-*dn*-*cargo*-*positive* embryos as scored by Cerulean fluorescence, Figure 6F) in line with mosaicism for *fgfr1*-*dn*-*cargo* activity. Besides activity of the used CreERT2 driver, the recombination efficiency of Tol2-based *loxP* transgenes is highly position-dependent (Carney and Mosimann, 2018); while our used transgenic insertion is functional, *de novo* generation of similar or even more potent *cargo* lines requires considerable screening effort. Further, since our experiments applied heatshock treatments relatively soon after CreERT2-mediated priming, future characterization is warranted to define if primed *fgfr1*-*dn*-*cargo* remains silent during prolonged phases without heatshock after priming, and how the primed transgene behaves upon repeated heatshock treatments in long-term experiments. Lastly, extension of this transgenic approach requires generation and validation of functional *lox* cassette excision of the desired cargo transgene insertion, which is notoriously sensitive to position effects and requires extensive screening for functional lines (Carney and Mosimann, 2018; Felker and Mosimann, 2016). Altogether, the *fgfr1*-*dn*-*cargo* line provides a transgenic tool to precisely perturb the FGF pathway during developmental processes based on the paired CreERT2 driver.

## Materials and Methods

### Zebrafish husbandry

Wildtype and transgenic zebrafish were raised and maintained at 28.5°C without light cycle essentially as described (Westerfield, 2007) and in agreement with procedures mandated by UZH and the veterinary office of the Canton of Zürich. Embryos were kept in E3 medium and strictly staged according to morphological characteristics corresponding to hours post-fertilization (hpf) or days post-fertilization (dpf) as described previously (Kimmel et al., 1995).

### Vectors and transgenic lines

Cloning reactions to create transgenesis vectors were performed with the Multisite Gateway system with LR Clonase II Plus (Life Technologies) according to the manufacturer’s instructions. The *fgfr1*-*dn*-*cargo* plasmid (*pAF019* or *pDestTol2CY*_*hsp70l:loxP*-*STOP*-*loxP*-*fgfr1a*-*dn*-*2A*-*Cerulean*-*CAAX*, *alpha*-*crystallin:YFP)* transgene was assembled from *pDH083* (Hesselson et al., 2009) by transfer of the *loxP* cassette into *pENTR5*’ (generating *pENTR/5*’_*hsp70l:loxP*-*STOP*-*loxP)*, with pAF017 (*pME*-*fgfr1a*-*dn*), and pAF018 (*p3E*-*2A*-*Cerulean*-*CAAX*) and *pCM326* (Mosimann et al., 2015) as backbone.

25 ng/μL Tol2 mRNA were injected with 25 ng/μL plasmid DNA for Tol2-mediated zebrafish transgenesis (Felker and Mosimann, 2016; Kwan et al., 2007). F0 founders were screened for specific *alpha*-*crystallin*:YFP expression, raised to adulthood, and screened for germline transmission. Single-insertion transgenic strains were established and verified through screening for a 50% germline transmission rate in outcrosses in subsequent generations as per our previously outlined procedures (Felker and Mosimann, 2016). We screened *α*-*crystallin* :YFP-expressing Tol2-generated F0 founders for functional transgene expression upon *cre* mRNA injection-based *loxP* excision followed by heatshock-mediated transgene activation to observe Cerulean-CAAX fluorescence and possible phenotypes; over a dozen founders with independent insertions needed to be screened to recover one functional transgenic line.

Previously established transgenic zebrafish lines used for this study include *ubi:creERT2* (expressing *myl7:EGFP* as transgenic marker) (Mosimann et al., 2011) and *drl:creERT2* (expressing *α*-*crystallin:Venus* as transgenic marker) (Mosimann et al., 2015).

### CreERT2/loxP experiments

*ubi:creERT2* or *drl:creERT2* transgenic zebrafish were individually crossed to the *fgfr1*-*dn*-*cargo* line. Embryos were induced using 4-OHT (Sigma H7904) from fresh and/or pre-heated (65° C for 10 minutes) stock solutions in DMSO with a final concentration of 10 μM in E3 embryo medium as per our established protocols (Felker and Mosimann, 2016; Felker et al., 2016). Heatshock treatments were performed for 1 h in E3-filled glass tubes in a 37°C water bath (measured and calibrated with thermometer in the water bath) at specific developmental stages as indicated in individual experiments. Double-transgenic embryos were detected though Cerulean-CAAX expression after heatshock using standard microscopy.

### Chemical treatments

Wildtype embryos were treated with SU5402 to globally perturb FGF signaling at the respective developmental stage. Single-use 100 mM SU5402 stock aliquots were thawed and diluted in E3 to a working concentration indicated in individual experiments directly before administration to the embryos. For pulsed SU5402-treatments, embryos were washed several times in fresh E3 medium after the desired incubation periods. Of note, we observed decreasing potency of stored SU5402 aliquots over time, warranting the use of fresh compound for experiments that require maximal FGF signaling inhibition (data not shown).

### Genomic DNA isolation and genotyping

Genomic DNA was isolated by incubating single embryos in 50 μL of 50mM NaOH at 95°C for 30 minutes and subsequent neutralization with 5 μl 1M Tris-HCl buffer (pH 5.0). Samples were spun down to remove debris and stored at 4°C until further use.

For genotyping potential *creERT2* transgene carriers, *oAF089* (*5*′-*GCATTACCGGTCGATGCAACG*-*3′)* and *oAF090* (*5*′-*CCAGAGACGGAAATCCATCGC*-*3′)* primers were used. Primers flanking both *loxP* sites (*oAF040 5*′-*CGTCGACTCTAGAGGATCACG*-*3′* and *attB1*_*rev 5*′-*AGCCTGCTTTTTTGTACAAACTTG*-*3′)* were used to genotype for the *fgfr1*-*dn*-*cargo* transgene, enabling assessment of successful recombination, as the PCR yields a shorter product (166 bp) after excision of the *loxP*-flanked STOP cassette compared to the non-recombined transgene yielding a long product (1129 bp). Standard Go Taq^®^ Green Master Mix (Promega) conditions were used for PCR.

### In situ probe synthesis

Antisense RNA probes were designed for the genes *etv4/pea3*, *dusp6/Mkp3*, and *spry4.* First-strand complementary DNA (cDNA) was generated from wildtype zebrafish RNA isolated from different developmental stages using Superscript III First-Strand Synthesis kit (Invitrogen) and subsequently pooled (Mosimann et al., 2015). DNA templates were generated using first-strand cDNA as PCR template and following primers: i) *etv4* with *oAF169* (*5*’-*TTACGTATGCAGCCTTCTCG*-*3*’) and *oAF170* (*5*’-*GGTTCATGGGGTAACTGTGG*-*3*’); ii) *dusp6* with *oLK005* (*5*′-*CTCCGTGTTGGGTTTACTGC*-*3*′) and *oLK006* (*5*′-*AAGTACAGTGGCTGGGTTGG*-*3*′); *spry4* with *oAF183* (*5*’-*ACTGATGAGGACGAGGAAGG*-*3*’) and *oAF184* (*5*’-*GACTCGGAATCCTTCAGTGG*-*3*’). For *in vitro* transcription initiation, the T7 RNA polymerase promoter *5*′-*TAATACGACTCACTATAGGG*-*3*’ was added to the 5’-end of reverse primers. PCR reactions were performed under standard conditions using Phusion High-Fidelity DNA Polymerase (ThermoFisher Scientific). RNA probes were generated via overnight incubation at 37°C using T7 RNA polymerase (20 U/μl) (Roche) and Digoxigenin (DIG)-labeled dNTPs (Roche). The resulting RNA was treated with 1μM DNAse (Roche) for 15 minutes at 37°C and cleaned-up in lithium chloride and ethanol to precipitate the RNA.

### Embryo fixation and whole-mount in situ hybridization

Embryos were fixed in 4% Paraformaldehyde (PFA) overnight at 4°C, transferred into 100% methanol and stored at −20°C until *in situ* hybridization. *In situ* hybridization of whole-mount zebrafish embryos was performed according to published protocols (Thisse and Thisse, 2008).

### Microscopy and image analysis

Brightfield (BF), basic fluorescence, and *in situ* hybridization imaging was performed using a Leica M205FA equipped with a DFC450 C camera.

*In toto* fluorescent embryo imaging was performed by Single Plane Illumination Microscopy (SPIM) with a Zeiss Lightsheet Z.1 microscope. Prior to imaging, embryos were embedded in a rod of 1% low melting agarose in E3 with 0.016% Ethyl 3-aminobenzoate methanesulfonate salt (Tricaine, Sigma) in a 50 μL glass capillary. During acquisition, embedded zebrafish were kept at 28°C in a chamber containing E3 with 0.016% Tricaine. Imaging was performed from three to four angles and images from all illumination sources were fused using the Zeiss Zen Black software. Zeiss Zen Black was also used to construct maximum intensity projections.

All further image processing with Leica LAS, ImageJ/Fiji, Imaris and Adobe Photoshop and Illustrator CS6 according to image-preserving guidelines to ensure unbiased editing of the acquired image data.

### Statistics

All statistical analysis was performed using GraphPad Prism 6.0. Data are presented as mean ± SEM, if not noted otherwise. A lower case “n” denotes the number of embryos, while a capital “N” signifies the number of replicates. For comparison of two groups, a 2-tailed unpaired Student’s t-test was performed. A p-value of 0.05 was considered significant (p > 0.05 ns, p ≤ 0.05^∗^).

## Acknowledgements

We thank Sibylle Burger and Seraina Bötschi for technical and husbandry support, Dr. Stephan Neuhauss for zebrafish support, Dr. Daniela Panáková for assistance with phenotype analysis, and all members of the Mosimann lab for critical input.

## Author contributions

L.K., A.F., E.C., and C.M. designed, performed, and analyzed the experiments; L.K., A.F., and C.M. compiled data and wrote the manuscript.

## Supplementary Figure legends

**Supplementary Figure 1:**
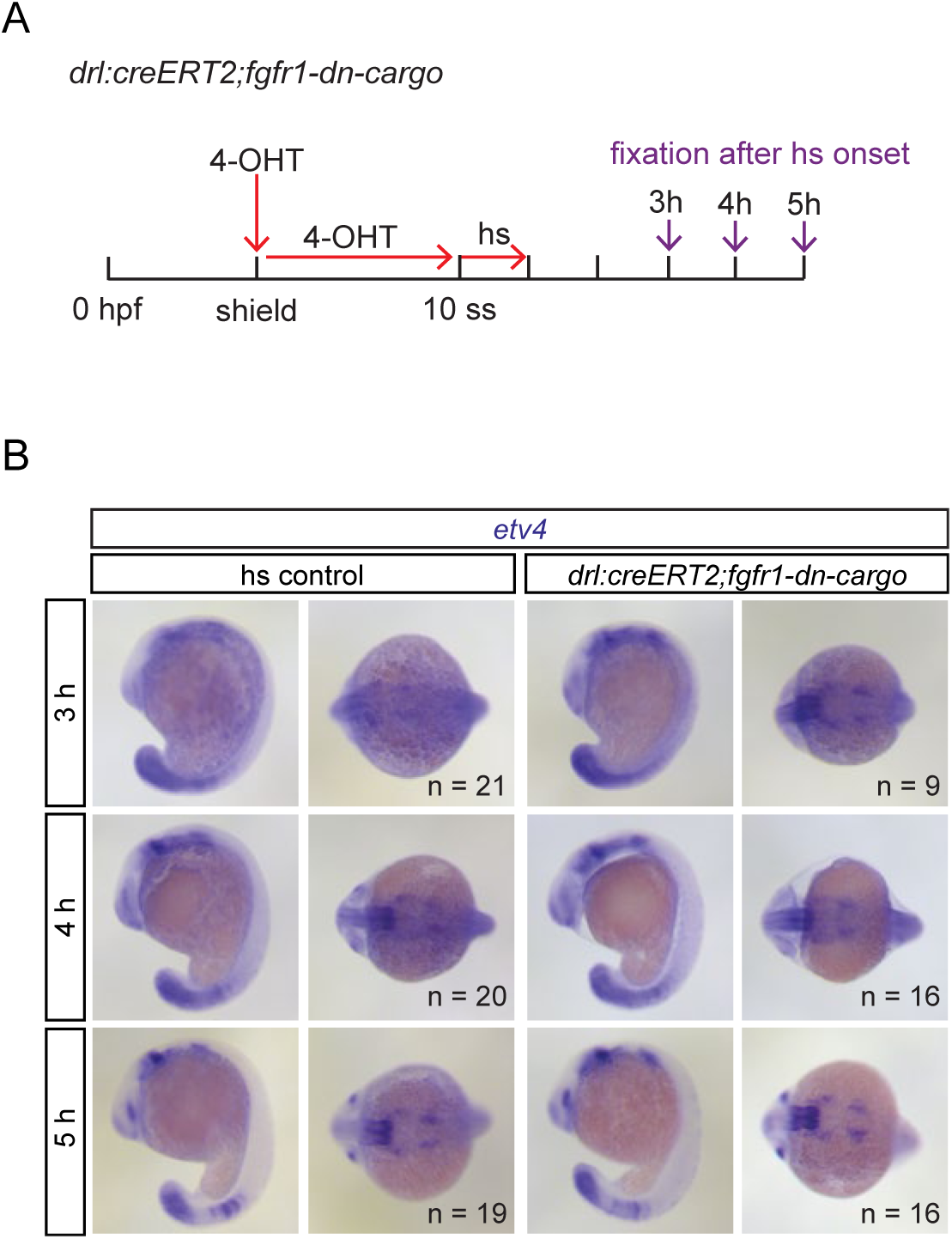
Embryo-wide FGF target gene expression remains grossly intact upon LPM-specific FGF signaling perturbation. (**A**) Schematic showing the timeline of treatments for genetic FGF signaling perturbations in *drl:creERT2;fgfr1*-*dn*-*cargo* and timepoints of embryo fixation for signaling activity read-outs. Red arrows indicate timepoints and duration of 4-OHT and heatshock treatments, violet arrows mark timepoints of embryo fixation after initiation of transgene activation via heatshock. (**B**) Embryos selectively perturbed for FGF signaling in the *drl* descendants at somitogenesis displayed no detectable differences in *etv4* expression when compared to unperturbed siblings 3-5 hours after heatshock treatment (lateral and dorsal views, anterior to the left). n indicates numbers of individual embryos analyzed for each condition.

**Supplementary Figure 2:**
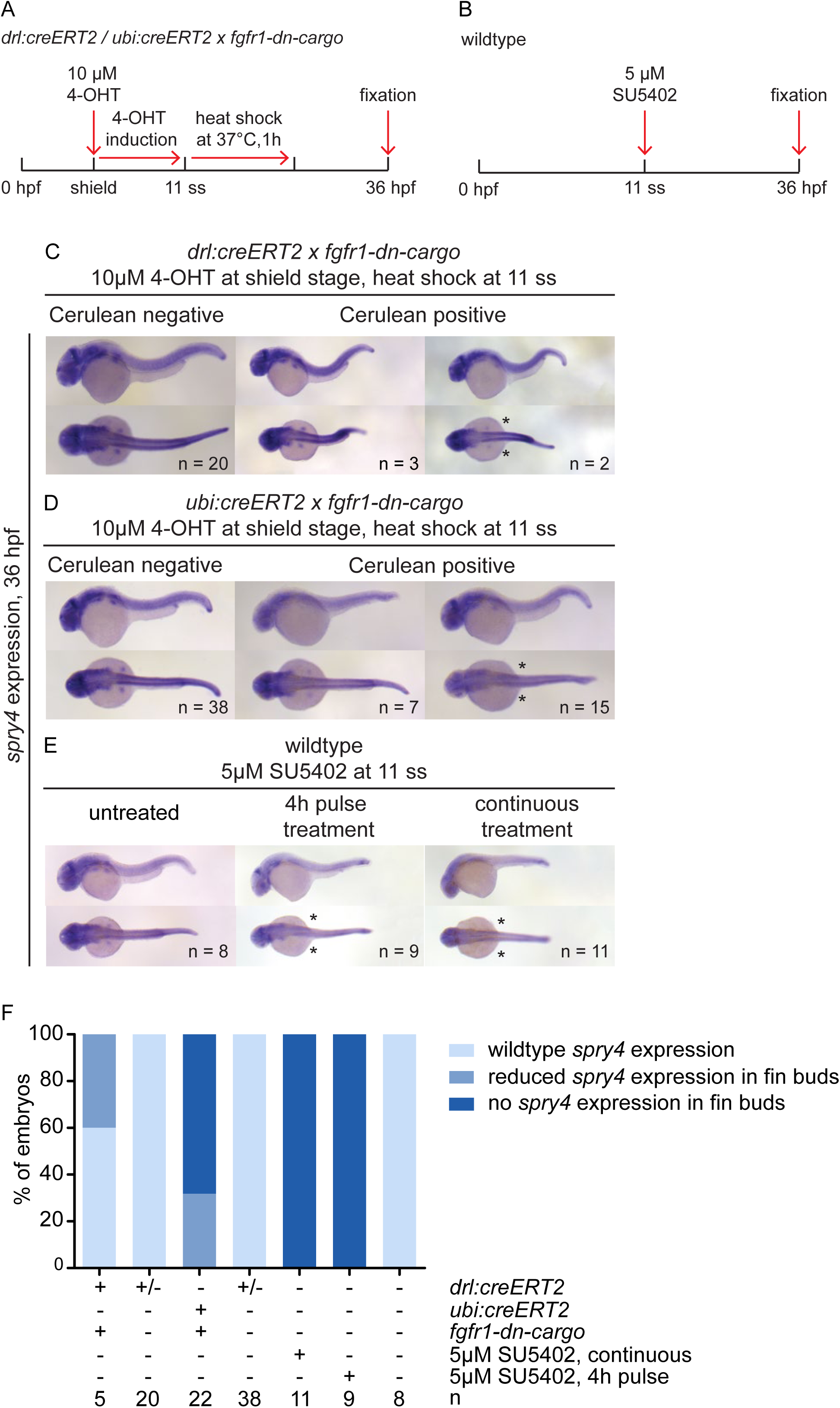
*spry4* expression in tissue-specifically and globally FGF signaling-perturbed embryos at 36 hpf. (**A, B**) Transgenic embryos were treated with 4-OHT at shield stage and heatshock-treated at 11 ss. Wildtype embryos were treated with SU5402 either continuously or for a four-hour pulse. (**C-E**) *spry4* expression was completely absent in embryos continuously treated with SU5402, and reduced or absent in fin buds of transgenic (*drl*-based LPM priming or *ubi*-based ubiquitous priming) or pulse-treated embryos; lateral and dorsal views, anterior to the left, asterisks indicate loss of pectoral fin expression. (**F**) Quantification of *spry4* expression in embryos subjected to embryo-wide or tissue-specific signaling perturbations. n indicates numbers of individual embryos analyzed for each condition.

**Supplementary Figure 3:**
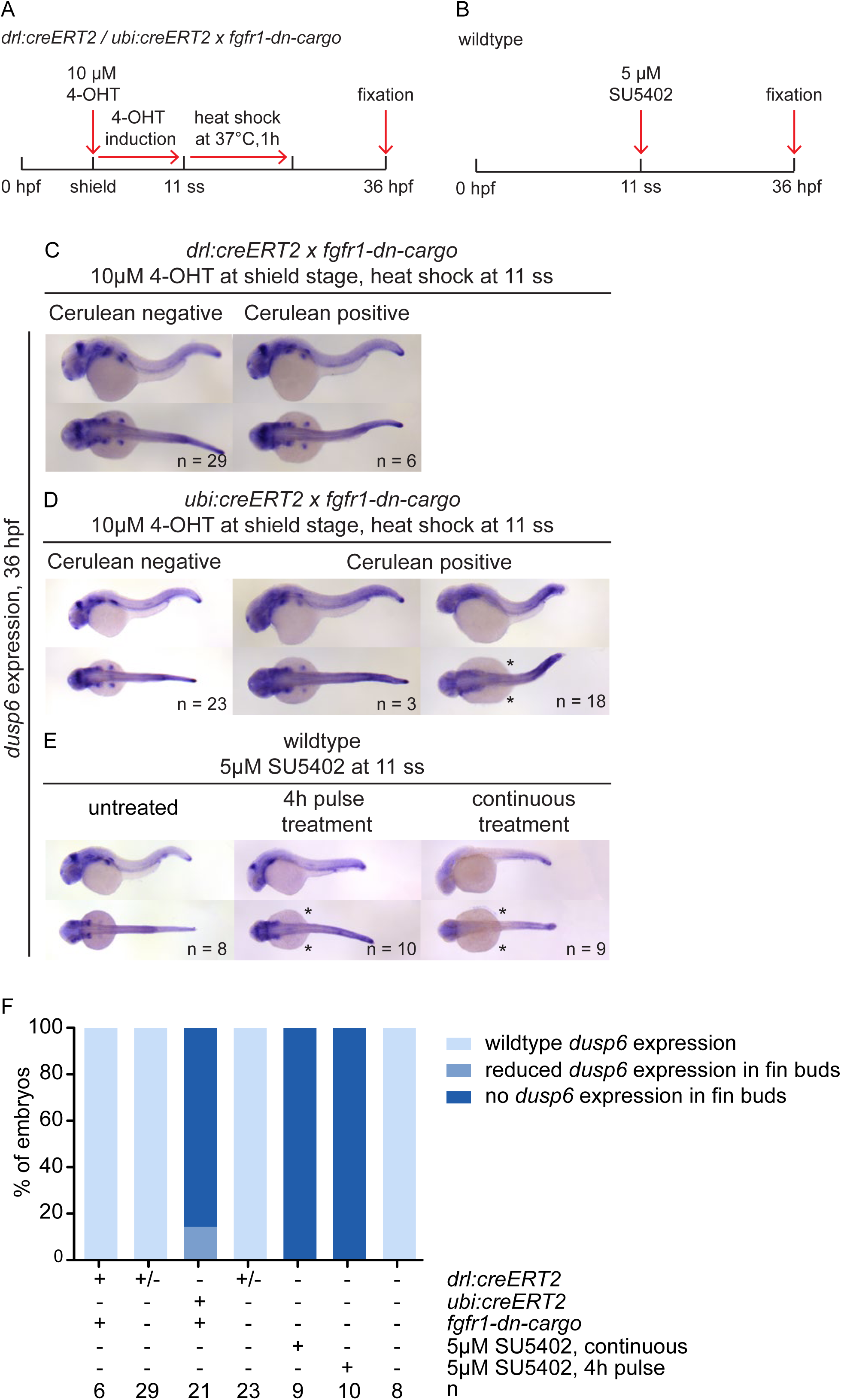
*dusp6* expression in tissue-specifically and globally FGF signaling-perturbed embryos at 36 hpf. (**A, B**) Transgenic embryos were treated with 4-OHT at shield stage and heatshock-treated at 11 ss. Wildtype embryos were treated with SU5402 either continuously or for a four-hour pulse. (**C-E**) *dusp6* expression was completely absent in embryos continuously treated with SU5402, and reduced or absent in fin buds of transgenic (*drl*-based LPM priming or *ubi*-based ubiquitous priming) or pulse-treated embryos; lateral and dorsal views, anterior to the left, asterisks indicate loss of pectoral fin expression. Note that *drl:creERT2*-primed embryos retained *dusp6* expression in pectoral fin fields, while *ubi:creERT2*-primed, ubiquitous *fgfr1a*-*dn* expression caused pectoral fin-specific loss in a cohort of embryos. (**F**) Quantification of *dusp6* expression in embryos subjected to global or tissue-specific signaling perturbations. n indicates numbers of individual embryos analyzed for each condition.

## References

Amaya, E., Musci, T. J. and Kirschner, M. W. (1991). Expression of a dominant negative mutant of the FGF receptor disrupts mesoderm formation in Xenopus embryos. Cell 66, 257–70.

Bökel, C. and Brand, M. (2013). Generation and interpretation of FGF morphogen gradients in vertebrates. Curr. Opin. Genet. Dev. 23, 415–422.

Böttcher, R. T. and Niehrs, C. (2005). Fibroblast growth factor signaling during early vertebrate development. Endocr. Rev. 26, 63–77.

Brand, M., Heisenberg, C. P., Jiang, Y. J., Beuchle, D., Lun, K., Furutani-Seiki, M., Granato, M., Haffter, P., Hammerschmidt, M., Kane, D. a, et al. (1996). Mutations in zebrafish genes affecting the formation of the boundary between midbrain and hindbrain. Development 123, 179–190.

Branda, C. S. and Dymecki, S. M. (2004). Talking about a revolution: The impact of site-specific recombinases on genetic analyses in mice. Dev Cell 6, 7–28.

Carney, T. J. T. J. and Mosimann, C. (2018). Switch and Trace: Recombinase Genetics in Zebrafish. Trends Genet. 0,.

Chen, J., Xia, L., Bruchas, M. R. and Solnica-Krezel, L. (2017). Imaging early embryonic calcium activity with GCaMP6s transgenic zebrafish. Dev. Biol.

Crossley, P. H., Minowada, G., Macarthur, C. A. and Martin, G. R. (1996). Roles for FGF8 in the Induction, Initiation, and Maintenance of Chick Limb Development. 84, 127–136.

de Pater, E., Clijsters, L., Marques, S. R., Lin, Y.-F. F., Garavito-Aguilar, Z. V, Yelon, D. and Bakkers, J. (2009). Distinct phases of cardiomyocyte differentiation regulate growth of the zebrafish heart. Development 136, 1633–1641.

Deng, C. X., Wynshaw-Boris, A., Shen, M. M., Daugherty, C., Ornitz, D. M. and Leder, P. (1994). Murine FGFR-1 is required for early postimplantation growth and axial organization. Genes Dev. 8, 3045–57.

Dong, P. D. S., Munson, C. A., Norton, W., Crosnier, C., Pan, X., Gong, Z., Neumann, C. J. and Stainier, D. Y. R. (2007). Fgf10 regulates hepatopancreatic ductal system patterning and differentiation. Nat. Genet. 39, 397–402.

Fallon, J. F., López, A., Ros, M. A., Savage, M. P., Olwin, B. B. and Simandl, B. K. (1994). FGF-2: apical ectodermal ridge growth signal for chick limb development. Science 264, 104–7.

Felker, A. and Mosimann, C. (2016). Contemporary zebrafish transgenesis with Tol2 and application for Cre/lox recombination experiments. Methods Cell Biol. 135, 219–44.

Felker, A., Nieuwenhuize, S., Dolbois, A., Blazkova, K., Hess, C., Low, L. W. L. W. L., Burger, S., Samson, N., Carney, T. J. T. J., Bartunek, P., et al. (2016). In Vivo Performance and Properties of Tamoxifen Metabolites for CreERT2 Control. PLoS One 11, e0152989.

Fischer, S., Draper, B. W. and Neumann, C. J. (2003). The zebrafish fgf24 mutant identifies an additional level of Fgf signaling involved in vertebrate forelimb initiation. Development 130, 3515–3524.

Hans, S., Kaslin, J., Freudenreich, D. and Brand, M. (2009). Temporally-controlled site-specific recombination in zebrafish. PLoS One 4, e4640.

Hans, S., Freudenreich, D., Geffarth, M., Kaslin, J., Machate, A. and Brand, M. (2011). Generation of a non-leaky heat shock-inducible Cre line for conditional Cre/lox strategies in zebrafish. Dev. Dyn. 240, 108–15.

Henninger, J., Santoso, B., Hans, S., Durand, E., Moore, J., Mosimann, C., Brand, M., Traver, D. and Zon, L. (2017). Clonal fate mapping quantifies the number of haematopoietic stem cells that arise during development. Nat. Cell Biol. 19,.

Hess, C., Prummel, K. D., Nieuwenhuize, S., Parker, H., Rogers, K. W., Kozmikova, I., Racioppi, C., Burger, S., Brombacher, E. C., Burger, A., et al. (2018). A conserved regulatory program drives emergence of the lateral plate mesoderm. bioRxiv 261115.

Hesselson, D., Anderson, R. M., Beinat, M. and Stainier, D. Y. (2009). Distinct populations of quiescent and proliferative pancreatic beta-cells identified by HOTcre mediated labeling. Proc Natl Acad Sci U S A 106, 14896–14901.

Huang, P. and Stern, M. J. (2005). FGF signaling in flies and worms: More and more relevant to vertebrate biology. Cytokine Growth Factor Rev. 16, 151–158.

Itoh, N. and Konishi, M. (2007). The zebrafish fgf family. Zebrafish 4, 179–186.

Kimmel, C. B., Ballard, W. W., Kimmel, S. R., Ullmann, B. and Schilling, T. F. (1995). Stages of embryonic development of the zebrafish. Dev Dyn 203, 253–310.

Kroll, K. L. and Amaya, E. (1996). Transgenic Xenopus embryos from sperm nuclear transplantations reveal FGF signaling requirements during gastrulation. Development 122, 3173–83.

Kwan, K. M., Fujimoto, E., Grabher, C., Mangum, B. D., Hardy, M. E., Campbell, D. S., Parant, J. M., Yost, H. J., Kanki, J. P. and Chien, C. Bin (2007). The Tol2kit: a multisite gateway-based construction kit for Tol2 transposon transgenesis constructs. Dev Dyn 236, 3088–3099.

Ledda, F. and Paratcha, G. (2007). Negative regulation of Receptor Tyrosine Kinase (RTK) signaling: a developing field. Biomark. Insights 2, 45–58.

Lee, Y., Grill, S., Sanchez, A., Murphy-Ryan, M. and Poss, K. D. (2005). Fgf signaling instructs position-dependent growth rate during zebrafish fin regeneration. Development 132, 5173–83.

Lewandoski, M., Sun, X. and Martin, G. R. (2000). Fgf8 signalling from the AER is essential for normal limb development. Nat. Genet. 26, 460–463.

Mariani, F. V., Ahn, C. P. and Martin, G. R. (2008). Genetic evidence that FGFs have an instructive role in limb proximal–distal patterning. Nature 453, 401–405.

Marques, S. R., Lee, Y., Poss, K. D. and Yelon, D. (2008). Reiterative roles for FGF signaling in the establishment of size and proportion of the zebrafish heart. 321, 397–406.

Mercader, N., Fischer, S. and Neumann, C. J. (2006). Prdm1 acts downstream of a sequential RA, Wnt and Fgf signaling cascade during zebrafish forelimb induction. Development 133, 2805–2815.

Mohammadi, M., McMahon, G., Sun, L., Tang, C., Hirth, P., Yeh, B. K., Hubbard, S. R. and Schlessinger, J. (1997). Structures of the Tyrosine Kinase Domain of Fibroblast Growth Factor Receptor in Complex with Inhibitors. Science (80-.). 276,.

Moon, A. M. and Capecchi, M. R. (2000). Fgf8 is required for outgrowth and patterning of the limbs. Nat. Genet. 26, 455–459.

Mosimann, C. and Zon, L. I. (2011). Advanced zebrafish transgenesis with Tol2 and application for Cre/lox recombination experiments. Methods Cell Biol 104, 173–194.

Mosimann, C., Kaufman, C. K. C. K., Li, P., Pugach, E. K. E. K., Tamplin, O. J. O. J. and Zon, L. I. L. I. (2011). Ubiquitous transgene expression and Cre-based recombination driven by the ubiquitin promoter in zebrafish. Development 138, 169–177.

Mosimann, C., Panáková, D., Werdich, A. A. A., Musso, G., Burger, A., Lawson, K. L. K. L., Carr, L. A. L. A., Nevis, K. R. K. R., Sabeh, M. K. K., Zhou, Y., et al. (2015). Chamber identity programs drive early functional partitioning of the heart. Nat. Commun. 6, 8146.

Niswander, L., Tickle, C., Vogel, A., Booth, I. and Martin, G. R. (1993). FGF-4 replaces the apical ectodermal ridge and directs outgrowth and patterning of the limb. Cell 75, 579–87.

Nomura, R., Kamei, E., Hotta, Y., Konishi, M., Miyake, A. and Itoh, N. (2006). Fgf16 is essential for pectoral fin bud formation in zebrafish. Biochem. Biophys. Res. Commun. 347, 340–346.

Norton, W. H. J., Ledin, J., Grandel, H. and Neumann, C. J. (2005). HSPG synthesis by zebrafish Ext2 and Extl3 is required for Fgf10 signalling during limb development. Development 132, 4963–4973.

Ohuchi, H., Nakagawa, T., Yamamoto, A., Araga, A., Ohata, T., Ishimaru, Y., Yoshioka, H., Kuwana, T., Nohno, T., Yamasaki, M., et al. (1997). The mesenchymal factor, FGF10, initiates and maintains the outgrowth of the chick limb bud through interaction with FGF8, an apical ectodermal factor. Development 124, 2235–44.

Oki, S., Kitajima, K. and Meno, C. (2010). Dissecting the role of Fgf signaling during gastrulation and left-right axis formation in mouse embryos using chemical inhibitors. Dev. Dyn. 239, 1768–1778.

Ornitz, D. M. and Itoh, N. (2001). Protein family review: Fibroblast growth factors. Genome Biol. 2, reviews3005.1-3005.12.

Ornitz, D. M. and Itoh, N. (2015). The Fibroblast Growth Factor signaling pathway. Wiley Interdiscip. Rev. Dev. Biol. 4, 215–66.

Ota, S., Tonou-Fujimori, N. and Yamasu, K. (2009). The roles of the FGF signal in zebrafish embryos analyzed using constitutive activation and dominant-negative suppression of different FGF receptors. 126, 1–17.

Ota, S., Tonou-Fujimori, N., Tonou-Fujimori, N., Nakayama, Y., Ito, Y., Kawamura, A. and Yamasu, K. (2010). FGF receptor gene expression and its regulation by FGF signaling during early zebrafish development. genesis 48, 707–716.

Plotnikov, A. N., Hubbard, S. R., Schlessinger, J. and Mohammadi, M. (2000). Crystal structures of two FGF-FGFR complexes reveal the determinants of ligand-receptor specificity. Cell 101, 413–424.

Pradhan, A., Zeng, X.-X. I., Sidhwani, P., Marques, S. R., George, V., Targoff, K. L., Chi, N. C. and Yelon, D. (2017). FGF signaling enforces cardiac chamber identity in the developing ventricle. Development 144, 1328–1338.

Prykhozhij, S. V and Neumann, C. J. (2008). Distinct roles of Shh and Fgf signaling in regulating cell proliferation during zebrafish pectoral fin development. BMC Dev. Biol. 8, 91.

Raible, F. and Brand, M. (2001). Tight transcriptional control of the ETS domain factors Erm and Pea3 by Fgf signaling during early zebrafish development. Mech. Dev. 107, 105–17.

Reifers, F., Böhli, H., Walsh, E. C., Crossley, P. H., Stainier, D. Y. R. and Brand, M. (1998). Fgf8 is mutated in zebrafish acerebellar (ace) mutants and is required for maintenance of midbrain-hindbrain boundary development and somitogenesis. Development 125, 2381–2395.

Reifers, F., Walsh, E. C., Léger, S., Stainier, D. Y. R., Brand, M., Leger, S., Stainier, D. Y. R. and Brand, M. (2000). Induction and differentiation of the zebrafish heart requires fibroblast growth factor 8 (fgf8/acerebellar). Development 127, 225–235.

Roehl, H. and Nüsslein-Volhard, C. (2001). Zebrafish pea3 and erm are general targets of FGF8 signaling. Curr. Biol. 11, 503–507.

Rossant, J. and Nagy, A. (1995). Genome engineering: the new mouse genetics. Nat. Med. 1, 592–4.

Sekine, K., Ohuchi, H., Fujiwara, M., Yamasaki, M., Yoshizawa, T., Sato, T., Yagishita, N., Matsui, D., Koga, Y., Itoh, N., et al. (1999). Fgf10 is essential for limb and lung formation. Nat. Genet. 21, 138–141.

Simoes, F. C., Peterkin, T., Patient, R., Simões, F. C., Peterkin, T. and Patient, R. (2011). Fgf differentially controls cross-antagonism between cardiac and haemangioblast regulators. Development 138, 3235–3245.

Sun, X., Meyers, E. N., Lewandoski, M. and Martin, G. R. (1999). Targeted disruption of Fgf8 causes failure of cell migration in the gastrulating mouse embryo. Genes Dev. 13, 1834–46.

Thisse, C. and Thisse, B. (2008). High-resolution in situ hybridization to whole-mount zebrafish embryos. Nat. Protoc. 3, 59–69.

Ullrich, A. and Schlessinger, J. (1990). Signal transduction by receptors with tyrosine kinase activity. Cell 61, 203–212.

Vogel, A., Rodriguez, C. and Izpisúa-Belmonte, J. C. (1996). Involvement of FGF-8 in initiation, outgrowth and patterning of the vertebrate limb. Development 122, 1737–50.

Westerfield, M. (2007). The Zebrafish Book: a guide for the laboratory use of zebrafish (Danio rerio). 5th ed. Eugene: University of Oregon Press.

Xu, X., Weinstein, M., Li, C., Naski, M., Cohen, R. I., Ornitz, D. M., Leder, P. and Deng, C. (1998). Fibroblast growth factor receptor 2 (FGFR2)-mediated reciprocal regulation loop between FGF8 and FGF10 is essential for limb induction. Development 125, 753–65.

